# Programmed-DNA Elimination in the free-living nematodes *Mesorhabditis*

**DOI:** 10.1101/2022.03.19.484980

**Authors:** Carine Rey, Caroline Launay, Eva Wenger, Marie Delattre

## Abstract

In most species, elaborate programs exist to protect chromatin and maintain its integrity over cell cycles and generations. However some species systematically undergo excision and elimination of portions of their genome in somatic cells in a process called programmed-DNA elimination (PDE). PDE involves the elimination of mainly repeated elements but also protein-coding genes. PDE has been described in approximately 100 species from very distinct phyla, and more extensively in the parasitic nematodes *Ascaris* and in the unicellular Ciliates. In Ciliates, where PDE is pervasive, the underlying mechanisms have been studied and involve small RNA-guided heterochromatinization. In striking contrast, chromatin recognition and excision mechanisms remain mysterious in Metazoans, because the study species are not amenable to functional approaches. Above all, the function of such a mechanism, which has emerged repeatedly throughout evolution, is unknown. Answering these questions will provide significant insights into our understanding of chromatin regulation and genome stability.

We fortuitously discovered the phenomenon of PDE in all species of the free-living nematode genus *Mesorhabditis. Mesorhabditis*, which belong to the same family as *C. elegans*, have a small ∼150 Mb genome and offer many experimental advantages to start elucidating the elimination mechanisms in Metazoans. In this first study, we have used a combination of cytological observation and genomic approaches to describe PDE in *Mesorhabditis*. We found that the dynamics of chromosome fragmentation and loss are very similar to those described in *Ascaris*. Elimination occurs once in development, at the third embryonic cell division in all 5 presomatic blastomeres. Similar to other species, *Mesorhabditis* eliminate repeated elements but also about a hundred unique sequences. Most of the eliminated unique sequences are either pseudogenes or poorly conserved protein-coding genes. Our results raise the possibility that PDE has not been selected for a gene regulatory function in *Mesorhabditis* but rather mainly is a mechanism to irreversibly silence repeated elements in the soma.

## Introduction

In most eukaryotes, all cells of an organism have an identical genome, and many mechanisms ensure its integrity. However, for about a hundred species from 9 phyla, including mammals, Programmed-DNA Elimination (PDE) leads to the systematic elimination of specific DNA sequences in the soma (Dedukh and Krasikova, 2022; Wang and Davis, 2014). PDE is more pervasive in the phylum of Ciliates and has been well studied in a few species, leading to seminal discoveries on the role of small RNAs in genome rearrangement. In these unicellular organisms, both germline (MIC) and somatic (MAC) genomes coexist. When a new MAC is formed after massive amplification of the MIC, it also undergoes PDE of non-coding repeated sequences. Small RNA-mediated heterochromatin formation by PIWI-interacting RNAs (piRNAs) is required to trigger the excision of some sequences, which is performed by the recruitment of a domesticated transposase. The excision of the other remaining sequences is performed by yet uncharacterized mechanisms (Bracht et al., 2013; Rzeszutek et al., 2020).

In Metazoans, the mechanisms of PDE have not been identified, most likely because the study species are not amenable to functional approaches. Genome sequencing performed on a few species revealed that most of the eliminated sequences are repetitive: transposable elements and satellite tandem repeats (Dedukh and Krasikova, 2022; Wang and Davis, 2014). Some protein-coding genes involved in germline development are also eliminated, suggesting PDE is a drastic solution to prevent the expression of unwanted sequences in the soma (Drotos et al., 2022; Maurer-Alcalá and Nowacki, 2019). Despite this striking evolutionary convergence, the pattern of elimination varies between species. For instance, in some birds and insects an entire chromosome is eliminated (the germline restricted chromosome) (Torgasheva et al., 2019). In copepods, after elimination the retained fragments join to reform chromosomes; chromosome number is unchanged but somatic chromosomes are smaller than germline chromosomes (Grishanin, 2014). In *Ascaris* or lampreys, chromosome number increases after PDE (Timoshevskiy et al., 2016). In *Ascaris*, elimination occurs at once, whereas in lampreys it is progressive, over several cell cycles. The amount of eliminated sequence also varies largely, even between closely related species. For instance, *Ascaris suum* and *Parascaris univalens* eliminate 10% and 90% of their genomes, respectively (Wang et al., 2017). Among the most studied organisms are the parasitic nematodes from the *Ascaris* genus. In *Ascaris*, as in *C. elegans*, blastomere identity is specified very early on in embryos (Haag et al., 2018). PDE starts at the third embryonic cell cycle, as the five presomatic founder cells enter mitosis. During anaphase, DNA fragments are found detached from the mitotic spindle, and are thus excluded from the reforming nuclei at the end of mitosis (Niedermaier and Moritz, 2000). Interestingly, the centromeric histone CEN-H3 which uniformly decorates the holocentric chromosomes of nematodes, is absent from the unretained fragments, suggesting that local loss of CEN-H3 and hence loss of microtubule attachment, is a necessary step for fragment exclusion (Kang et al., 2016). How CEN-H3 is specifically lost on these fragments is unknown. Finally, new telomeres are added on the retained fragments, which form new mini-chromosomes (Wang et al., 2020). In *Parascaris univalens*, the germline cells have 2n=2 large chromosomes, but ∼60 small chromosomes are found in the nuclei of somatic cells (Wang et al., 2017).

Recent chromosome-scale genome reconstruction in *Ascaris* has revealed that the eliminated satellite-rich regions are present in the subtelomeric regions of all chromosomes (Wang et al., 2020). Hence, the breakage of chromosomes in early embryos is also accompanied by removal of all telomeres, and telomeres are progressively reassembled at chromosome breakpoints after a few cell cycles (Wang et al., 2020). A recent study on the nematode *Oscheius tipulae* (belonging to the same family as *C. elegans*) has revealed another example of telomere resetting after elimination of subtelomeric satellite sequences, although chromosomes are not fragmented nor reduced (Gonzalez de la Rosa et al., 2021). Systematic analysis of breakpoint sequences did not reveal any sequence signature in *Ascaris*. Though the eliminated regions are roughly the same in two independent blastomeres, in two different individuals of the same species, or even between closely related species, the exact breakpoint positions can vary within a zone of a few kb (Wang et al., 2012). It is unclear if these inexact breakpoints occur because DNA excision does not rely on a sequence-specific mechanism or because the breakpoints are chewed back after the excision.

Besides satellite repeats and transposable elements, PDE also targets some protein coding genes. ∼1000 genes are eliminated in the soma of *Ascaris*. As expected for genes that are eliminated in the soma, their expression is restricted to the germline or early embryos. Interestingly, 35% of these genes are conserved in all nematodes and Gene Ontology analysis revealed they are enriched in processes related to germline function (Wang et al., 2017). These results suggest the eliminated genes have an important function in the germline or early embryogenesis. Although they represent only 5% of the total number of genes, their elimination through PDE may have an important regulatory function. Around 50% of the eliminated genes also have a paralog in the *Ascaris* genome that is not eliminated, suggesting PDE may also have a role in gene dosage (Wang et al., 2017). Analysis performed in lampreys revealed the same categories and proportions for the unique sequences targeted by PDE (Smith et al., 2018).

Importantly, in *Ascaris*, the sequences targeted by PDE are not enriched in repressive histone marks, and small RNAs corresponding to these sequences have not been detected. Furthermore, PIWI proteins are absent from the *Ascaris* genome, suggesting a different mechanism for PDE than those described for Ciliates (Wang et al., 2012, 2017). However, Ciliates are also subjected to chromosome breakage without subsequent fusion (Rzeszutek et al., 2020), by a yet to be discovered mechanism-raising the exciting possibility that this mechanism shares similarities with PDE in *Ascaris*.

We fortuitously discovered that all species from the *Mesorhabditis* genus also undergo PDE. *Mesorhabditis* are free-living nematodes which belong to the same family as *C. elegans* (Rhabditidae) but have very distantly related to *Ascaris* (Haag et al., 2018). The phylogenetic distance of *Mesorhabditis* to *C. elegans* is as far as *Pristionchus pacificus* or *Oscheius tipulae* to *C. elegans* (two other well-studied nematode genera (Haag et al., 2018)). Within *Mesorhabditis*, some species have an asexual reproductive system, whether others are male/female sexual species (Grosmaire et al., 2019; Launay et al., 2020). We analyzed several *Mesorhabditis* species and systematically found the pattern of DNA fragmentation and loss previously described in *Ascaris*, irrespective of their reproductive system. Hence, the entire *Mesorhabditis* genus is probably subjected to PDE.

In this study, we used a combination of cytological observation, genomic and transcriptomic analysis to describe the dynamics of elimination and the nature of the eliminated sequences in the asexual species *M. belari* and the sexual species *M. spiculigera* (Launay et al., 2020). *Mesorhabditis* nematodes have a small ∼150 Mb genome, share many experimental advantages with the model species *C. elegans*, including the ability to inactivate genes by RNAi and to modify the genome with CRISPR/Cas9 (Blanc & al. in prep), which will open the way to functional approaches. *Mesorhabditis* will therefore offer the possibility to start identifying the mechanisms of PDE in Metazoans.

## Results

### Cytological evidence of programmed-DNA elimination in Mesorhabditis nematodes

In fixed embryos of *Mesorhabditis belari* stained with Hoechst dye, we noticed the presence of many DNA fragments in the cytoplasm. We foundthat all embryos from approximately the 8-cell to 32-cell stage contain cytoplasmic DNA fragments (Figure 1). To understand the origin of these fragments, we first characterized the cell lineage of *M. belari* embryos using DIC recordings and fixed specimens (Figure 1A). Although the order of cell division varied compared to *C. elegans*, the organization of blastomeres was very similar. We thus will use the same names for blastomeres as in *C. elegans* and other nematodes. As in the majority of nematodes, the first embryonic division is asymmetric in size. We thus defined the posterior pole as the position where the smallest cell (P1) is born. A series of asymmetric divisions in the P lineage finally gave rise to a very small cell, which is positioned very posteriorly and shows a dense aspect of chromatin. By analogy with *C. elegans*, we defined this smallest cell as the P4 germline founder cell.

**Figure 1:**
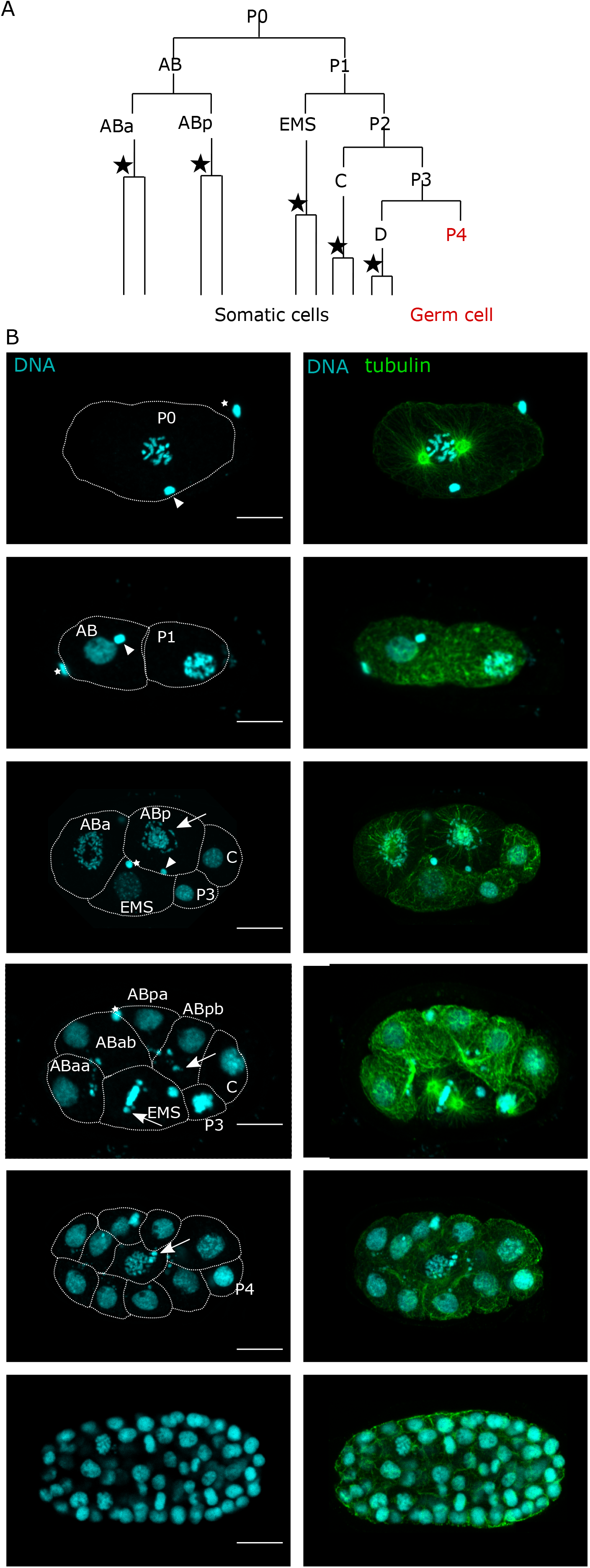
*M. belari* embryos eliminate portions of the genome in the 5 presomatic blastomeres. (A). Schematic representation of the early lineage of *M. belari* embryos. The lineage illustrates the order of division but not the exact timing between divisions. The black star represents a division of elimination. (B). Confocal images of *M. belari* embryos from 1-cell embryo (top) to ∼64-cell stage embryo (bottom). Unretained fragments (arrows) are clearly visible in 7-cell stage embryos, for instance in the cytoplasm of AB daughter cells or during the mitosis of EMS. Tubulin is in green, DNA in blue. White stars show the polar body. Scale bar is 10 um.

We found that cytoplasmic DNA fragments were first observed in the daughter cells of ABa and ABp (Figure 1B). Next, fragments were seen after the division of EMS. Fragments were also detected after the division of the C and the D blastomeres. Importantly, the P4 cell, which does not divide until hatching, did not contain any cytoplasmic fragments (Figure 1B). Once they appeared, the cytoplasmic fragments seemed to randomly segregate during the subsequent cell divisions. In *Ascaris* nematodes, the DNA fragments that are excluded from the reforming nuclei are visible for several cell cycles and are later destroyed, most likely by autophagy (Wang et al., 2020). Similarly, we found that the cytoplasmic DNA fragments are no longer visible after the ∼32 cell stage, suggesting they are also ultimately degraded (Figure 1B). The embryonic cell divisions of *M. belari* are slower than in *C. elegans* and we estimate that it is ∼6-8 hours between the time of production of cytoplasmic fragments and their ultimate elimination. Overall, the pattern of cytoplasmic fragment production and elimination strongly resembles what had been described in the parasitic nematodes *Ascaris*, suggesting that *M. belari* nematodes are also subjected to programmed-DNA elimination during their development.

We next asked if this phenomenon was restricted to *M. belari* within the *Mesorhabditis* genus. e analyzed the embryos of other *Mesorhabditis* species including asexual and sexual species (Launay et al., 2020). DNA staining of fixed embryos revealed the presence of cytoplasmic DNA fragments from the 3rd cell cycle onward, in all cells except the P4 cell (Figure S1), although the quantity of eliminated DNA seemed variable between species. We conclude that somatic programmed-DNA elimination is pervasive within the *Mesorhabditis* genus.

### Dynamics of DNA elimination in presomatic blastomeres

Our observation suggested that DNA elimination occurred during mitosis. To analyze the dynamics of fragment production and elimination, we imaged *M. belari* embryos at different time points during the cell cycle. We focused our analysis on the large cells ABa, ABp and EMS. Moreover, ABa and ABp divide perpendicular to the glass slide whereas EMS divides parallel to it. These orientations of division allowed us to obtain both transverse and longitudinal sections of the mitotic spindle.

M. belari has 2n=20 chromosomes. We found that before a division of elimination, at prometaphase, the number of chromosomes was higher and chromosomes appeared smaller (Figure 2). Hence, we hypothesize that chromosome breakage has occurred sometime between telophase and prometaphase. Second, we found that during the formation of the metaphase spindle, some of the chromosome fragments did not align on the metaphase plate. These fragments formed a halo around the metaphase plate (clearly visible in the ABa and ABp cells in a transversal view of the spindle), and seemed detached from the spindle microtubules (Figure 2). During anaphase, these fragments were not captured by the mitotic spindle and were not incorporated in the reforming nuclei (Figure 2).

**Figure 2:**
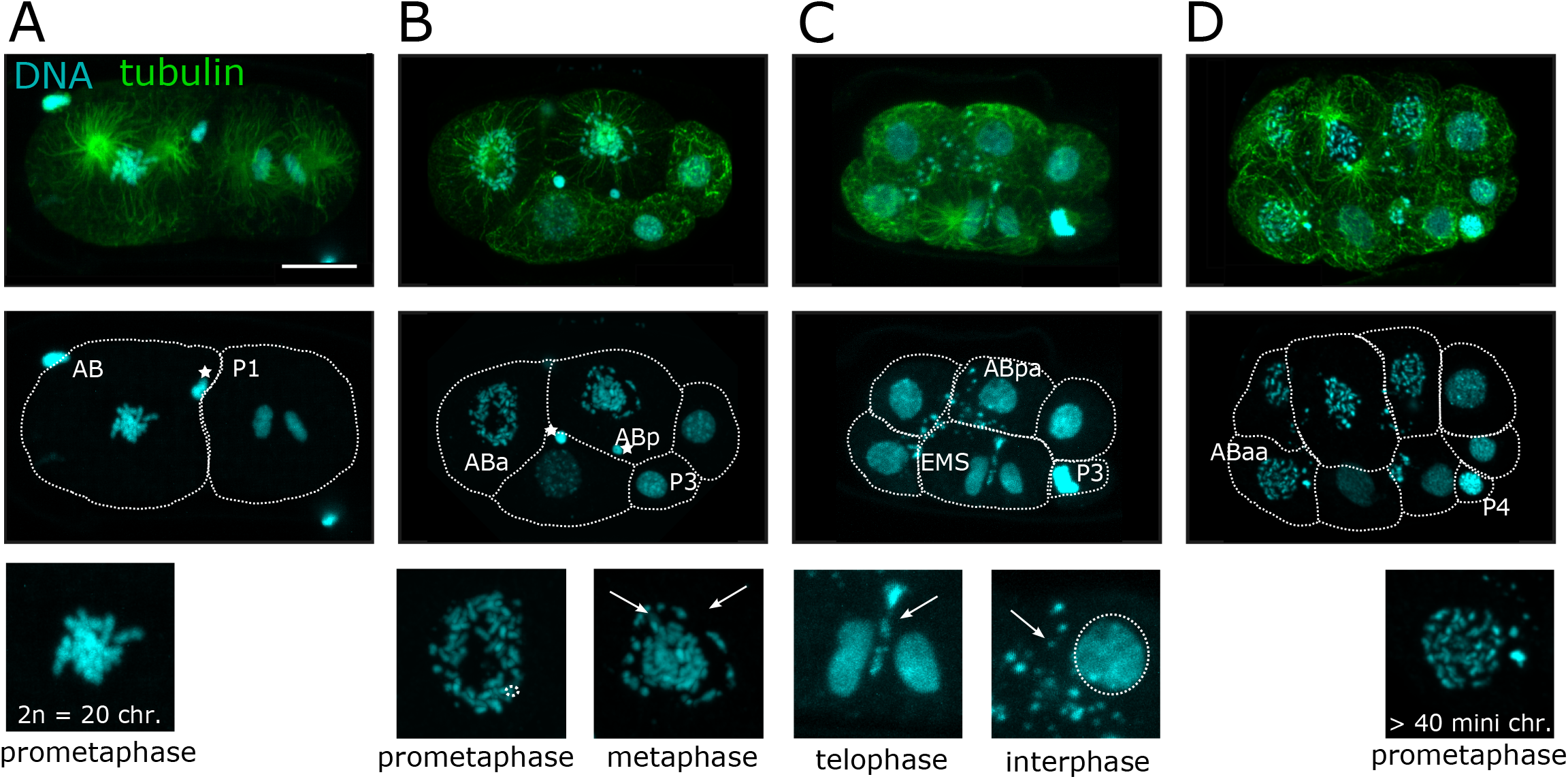
Dynamics of elimination during mitosis in *M. belari* embryos. One representative embryo per category is shown. Tubulin is in green and DNA is in blue. Scale bar is 10 um. Arrow points to the unretained fragments. (A). 2-cell stage embryo showing the 20 chromosomes in the AB nucleus. (B). 5-cell stage embryo. ABa and ABp are in metaphase (spindle oriented perpendicular to the glass-slide). Insets show the fragmented chromosomes and in the nuclei of ABp, some fragments form a halo around and detached from the metaphase plate. (C). 7-cell stage embryo during the division of EMS. The ABpp cell is not visible in this focal plane. EMS is in telophase and DNA fragments are excluded from the reforming nuclei. Many fragments are also visible outside the nuclei in ABpa. (D). 9-cell stage embryo. The daughters of ABa and ABp are in prometaphase and a minimum of 40 small chromosomes is counted. P3 and P4 cells are recognizable by their small size and highly compact DNA.

In *Ascaris*, fragments are seen lagging on the anaphase spindle during a division of elimination and those fragments are devoid of the centromeric histone CEN-H3. We used an antibody against the kinetochore protein BUB-1 and confirmed the loss of signal on the unretained fragments (Figure S2). This result suggests that, as in *Ascaris*, some fragments are excluded because they have lost the centromeric histones and cannot be captured by the spindle microtubules.

Importantly, during the subsequent cell divisions we did not detect new fragments that get excluded from the metaphase spindles, strongly suggesting that fragments are eliminated only once, as in *Ascaris*, during a single division of elimination, i.e at the third cell cycle (division of ABa, AB, EMS, C and D). Moreover, after a division of elimination, we counted ∼40 mini chromosomes, demonstrating that after elimination there was no fusion between the retained chromosomes and hence no restoration of the initial number of 20 chromosomes.

Overall, our results suggest that during a division of elimination, chromosomes are fragmented before or during prometaphase. At metaphase, some fragments lose their connection with the spindle microtubules (most likely due to the lack of centromeric proteins), and hence are lost in the cytoplasm where they are ultimately destroyed after a few cell cycles.

### Satellite tandem repeats and telomeres are eliminated in somatic cells

We next asked which sequences are eliminated in *M. belari* somatic cells. Eliminated sequences can be identified by comparing the genome of the germline (intact genome) and those of the soma (reduced genome), using DNAseq of dissected tissues. However, because of the very small size of *Mesorhabditis* nematodes (∼750 um), tissue dissection was technically challenging, and we had concerns about the amount of DNA and contamination between tissues. To circumvent these problems, we sequenced the genome of a pure population of L1 larvae, which are composed of ∼500 somatic cells and only 1 germline cell (see Mat and Methods). We also sequenced the genomes of adult females or males, which have a fully developed gonad and hence a much lower ratio of soma/germ cells compared to L1s. In particular, the large number of sperm cells in the gonad of males allowed us to obtain a higher proportion of the germline genome.

From the Illumina reads, we first analyzed the proportion of k-mers (i.e. subsequences of length k). We identified k-mers whose proportion was very high in adults compared to L1s, suggesting they are mainly present in the germline (Figure 3A). The most abundant k-mers corresponded to a small highly repeated sequence of 11nt (Sat11: 5’-GGAACCGTATT) and another one of 65nt (Sat65-5’-AGCTCAAATAGACTGAATAGCCTATTCTGAGCTGAATAGGCAATCTACACTTAATACCGTAATCC). To confirm that these satellite repeats are eliminated in the soma, we performed DNA FISH in early embryos using a fluorescent probe against each of these two satellites (Figure 3 and Figure S3A). We found a strong signal in all nuclei in early embryos, which was restricted to the cytoplasmic fragments as the cells entered a division of elimination. This result confirmed that elimination occurs during a single event and is not progressive over several cell cycles. In old embryos and in L1 larvae, the signal was detected in only 1 cell, which likely corresponded to the founder cell of the germline P4. P4 is also easily recognizable by its very small size and dense chromatin, as found in *C. elegans*. We concluded that these satellites are one target of programmed-DNA elimination in *M. belari*. We then used the same strategy, which does not require a fully assembled genome, to search for eliminated repeats in the genome of *M. spiculigera*. We performed DNAseq on L1s and adults and also uncovered germline-restricted satellite repeats (Figure S3B). Elimination of *M. spiculigera* Sat23 (5’AGGTGCAGGAGCCGAAACTATTT) was also confirmed by DNA FISH (Figure S3C). Hence, both species eliminate satellite repeats in their somatic cells, although these satellites have distinct sequences.

**Figure 3:**
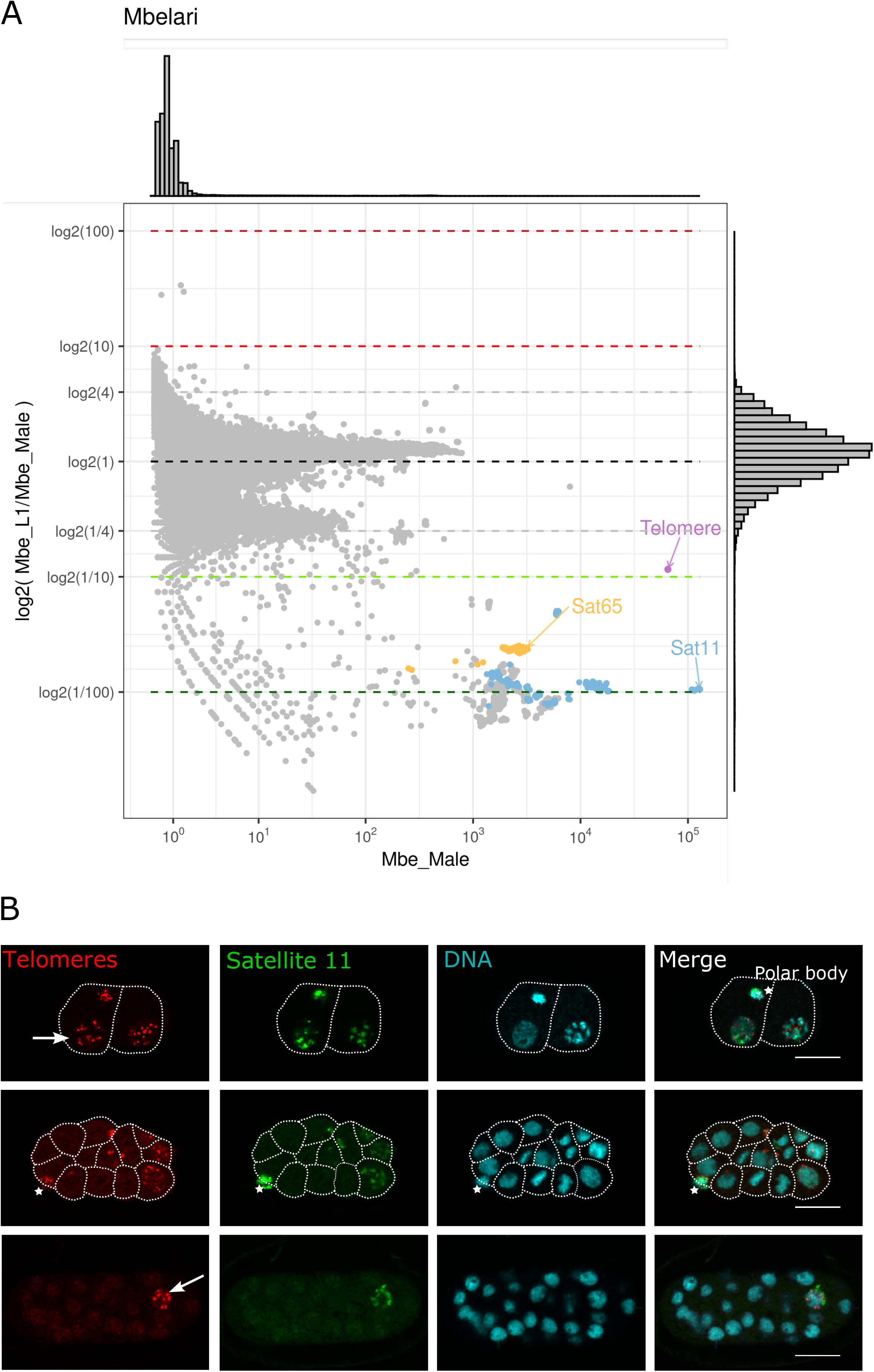
Identification of the eliminated satellite-tandem repeats. (A) k-mer analysis in L1 larvae, which contain mainly somatic tissues (and a single germline cell) and adult males, which contain a high number of germ cells (i.e. sperm cells). The ratio of counts between L1s and males is shown in log2 relative to the counts found in males. 1 grey dot represents one 56-mer. The majority of k-mers are unique and equally represented in both samples (log2(1)). K-mers containing the telomeric repeat are slightly overrepresented in males and overall are extremely abundant (10e5). Some k-mers, highlighted in colors, are underrepresented in L1s. K-mers shown in yellow and blue are different versions of the same 65nt and 11nt sequences, respectively. (B). DNA FISH validation of Sat11 (in green) and telomere (in red) elimination in *M. belari* embryos. DNA is in blue, scale bar is 10 um. Before elimination, the probes are detected in all nuclei of a 2-cell embryo (upper embryo). During elimination, the probes are only visible in the cells that have not yet entered a division of elimination and in the germline cell (middle embryo). In a ∼32-cell stage embryo, the probes are visible in a single nuclei, most likely the P4 germline cell. The white stars show the polar body.

In *Ascaris* nematodes, satellite repeats are also the target of elimination. Because some of these satellites are concentrated in subtelomeric regions, their elimination leads to telomere elimination in somatic cells (Wang et al., 2020). We thus tested if telomeres were also cleaved in young *M. belari* and *M. spiculigera* embryos. Our k-mer analysis identified the conserved telomeric repeats of nematodes 5’-TTACGG (Figure 3). Using a DNA FISH probe, we found that telomeres are eliminated at the same time as the satellite repeats in all somatic cells of *M. belari* and *M. spiculigera* embryos (Figure 3, Figure S3D), whereas a single cell, most likely the P4 germline cell, remained positive for the probe. To our surprise, we did not see the reappearance of telomeres in the soma of embryos or even L1 larvae, raising the possibility that embryonic cells continued cycling without telomeres. However, this result was inconsistent with our k-mer analysis which detected a lot of telomeric repeats in L1s (Figure 3). We hypothesize that as new telomeres reform in somatic nuclei, they add repeats slowly and are initially undetectable by DNA FISH.

Overall, our results show that similar to *Ascaris, Mesorhabditis* nematodes eliminate some satellite repeats in the genome of somatic nuclei, together with telomeric repeats, most likely because some of these satellites are present in subtelomeric regions.

### Transposable elements and some protein-coding genes are eliminated

We next asked if other types of sequence were eliminated in the somatic nuclei of *Mesorhabditis*. The analysis of k-mer missed the sequences that are not highly covered, i.e. unique sequences or imperfect repeats. We thus proceeded to map the DNAseq reads (windows of 100 bp, see Material and Methods) from *M. belari* L1s and adults onto the assembled genome (Grosmaire et al., 2019). We expected the eliminated sequences to have much fewer reads in L1s than in adults, whereas the retained fragments should have the same coverage in both samples. However, the ratio of coverage was not sufficiently distinct to retrieve the eliminated regions. We thus compared the ratio of coverage between L1s and males to the ratio of coverage between L1s and females. This approach allowed us to split the 100bp regions into 3 major clusters (Figure 4A). Most 100nt regions were found in the central cluster, i.e. similarly covered in all samples, suggesting these are the retained fragments. Some reads were biased towards the male sample and most likely corresponded to the Y chromosome. *M. belari* males are XY and hence the Y chromosome is absent in females, present in males and in 10% of L1s because the sex ratio is highly biased towards females in this species (Grosmaire et al., 2019). Some of the genes we had previously identified on the Y chromosome did map on this cluster, validating our clustering approach. The third cluster corresponded to reads that were overrepresented in both males and females compared to L1s, suggesting they were eliminated in L1s. Among those reads, we found Sat11 and Sat65, confirming that this cluster corresponded to the eliminated regions.

**Figure 4:**
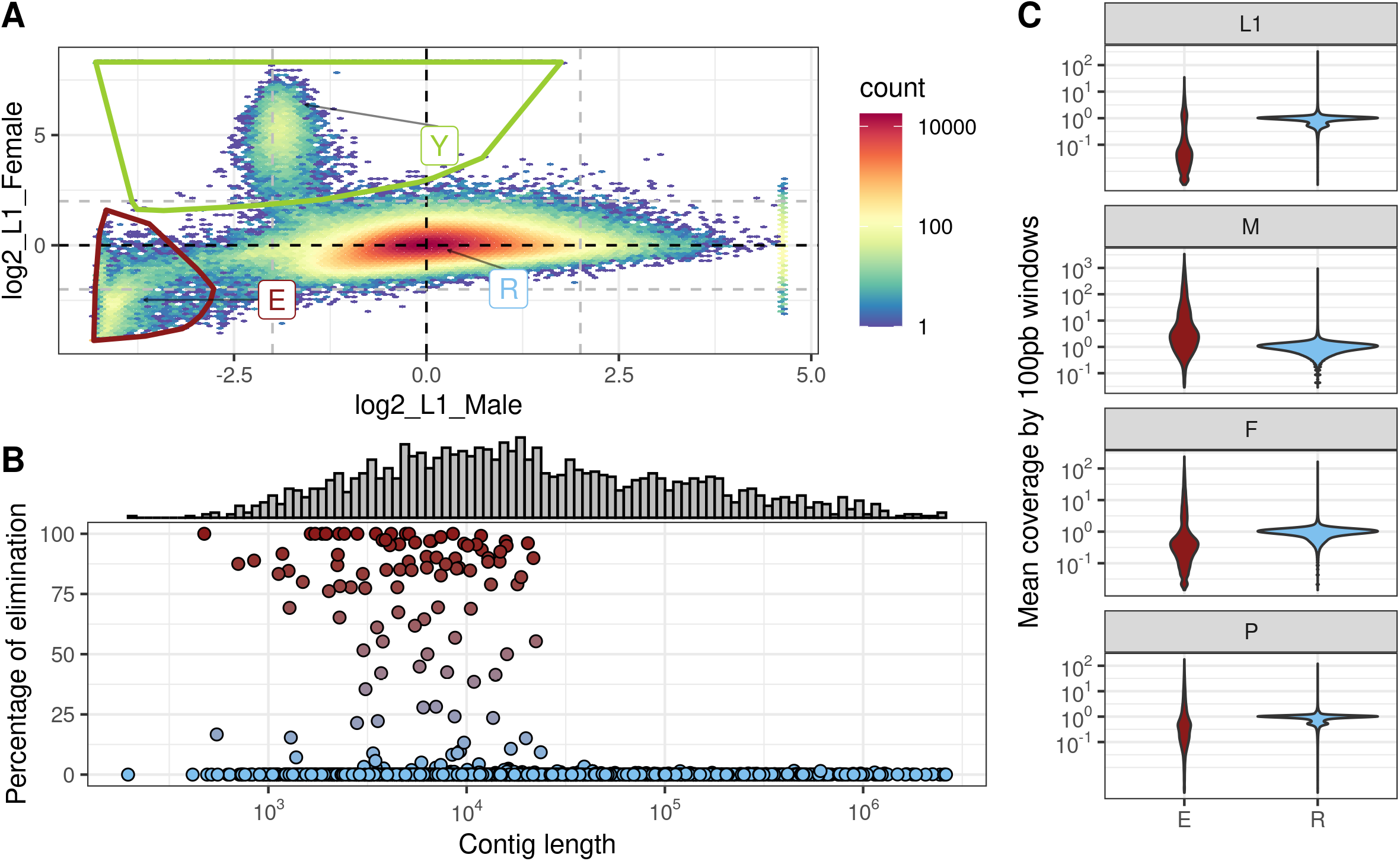
Clustering method to identify eliminated sequences in *M. belari*. (A). Analysis of coverage of 100nt-long segments from L1s (mainly soma), adult females or males (large proportion of germline) on the assembled genome. Ratio of coverage between L1s and females is shown in log2 relative to the ratio found between L1s and males. Cluster R likely corresponds to the retained DNA (germline restricted). A cluster of fragments overrepresented in males correspond to sequences from the Y chromosome of *M. belari* males. The E cluster likely represents the sequences that are eliminated, because they are underrepresented in L1s. (B). The assembled genome of *M. belari* is still highly fragmented in many contigs of variable length as shown on the histogram. For each contig (dot), the percentage of eliminated sequences is shown relative to the size of the contig in nt. A large number of small contigs (less than 10 kb) contain only eliminated sequences. (C). Validation of the normalization of read counts in the different samples, for the eliminated sequences (in red) and the retained sequences (in blue). In L1, the coverage of retained sequences is 1 and is below 1 for the eliminated sequences. As expected, the coverage of eliminated sequences is overrepresented in males for instance.

To further analyze these eliminated sequences, we mapped them back on the genome. We found that they were often found at the extremities of large contigs. Also, some contigs were entirely covered by these eliminated reads, showing that fragments of at least 10 kb can be entirely eliminated (Figure 4B). The current assembly of the genome is highly fragmented with many small contigs and our results show that many of these small contigs are eliminated (Figure 4B). These results suggested that i) the elimination of some genomic portions has perturbed the assembly of the genome, most likely because of the repeated nature of the eliminated fragments, ii) elimination mainly concerns large portions of the genome as previously found in *Ascaris*, rather than small sequences scattered in the genome, as found in Ciliates. As a final validation of our clustering approach and normalization of read counts, we confirmed that the normalized coverage of sequences from the retained cluster (in blue) was around one in all samples (Figure 4C). As expected, the coverage of reads from the eliminated cluster (in red) was very low in L1s (which contain mainly a reduced genome) but was much higher than 1 in females and even more in males (which contain a larger proportion of intact genome) (Figure 4C).

We next proceeded to analyze the nature of the eliminated sequences. Besides satellite repeats, we found that transposable elements (which we have not further analyzed in this study) and also 123 unique sequences were eliminated. To confirm these results, we randomly chose 6 of those eliminated genes and performed soma-specific PCR (Figure S4, see Material and Methods). Hence, as found in other Metazoans, genome elimination in *M. belari* targets repeated elements as well as unique sequences or genes.

### Most of the eliminated unique sequences are pseudogenes or are poorly conserved protein-coding genes

We next explored the nature and function of the eliminated genes. First, we analyzed their expression profiles. We analyzed read coverage in RNAseq data from a mixed population (all developmental stages, all tissues), only L1 larvae (i.e most likely only somatic expression), or in dissected adult male or female gonads. First, as expected from genes that are eliminated in the soma, none of these genes were expressed in L1s. Surprisingly, 63 out of the 123 genes were not expressed in any of the samples. Hence, it is very likely that these sequences correspond either to pseudogenes or intergenic sequences, which have been mis predicted as genes by the assembler. The remaining 60 sequences were true protein-coding genes, some being sex specific (Figure 5 and Table S1). Out of these 60 protein-coding genes, 38 had no ortholog in any species, and did not contain any predicted protein domains (Table S1). 11 protein-coding genes had an ortholog in *C. elegans*, among which only 2 genes were also germline-specific genes in *C. elegans (spe-49*, encoding a sperm membrane protein and *daz-1* encoding an RNA-binding protein). Nevertheless, most of these genes were highly expressed in the gonad of *M. belari*, in particular in the male gonad, suggesting they are *Mesorhabditis*-specific germline genes (Figure 5). Thus, out of the ∼1200 germline expressed genes identified in our comparative RNAseq approach, 60 genes were eliminated. Hence, 5% of the germline-specific genes are eliminated in the soma of *M. belari* during PDE. Overall, this is only 0.02% of all the protein-coding genes of *M. belari* (∼21000 genes (Grosmaire et al., 2019)) that are eliminated, which is much lower than the proportions found in *Ascaris* or lampreys.

**Figure 5:**
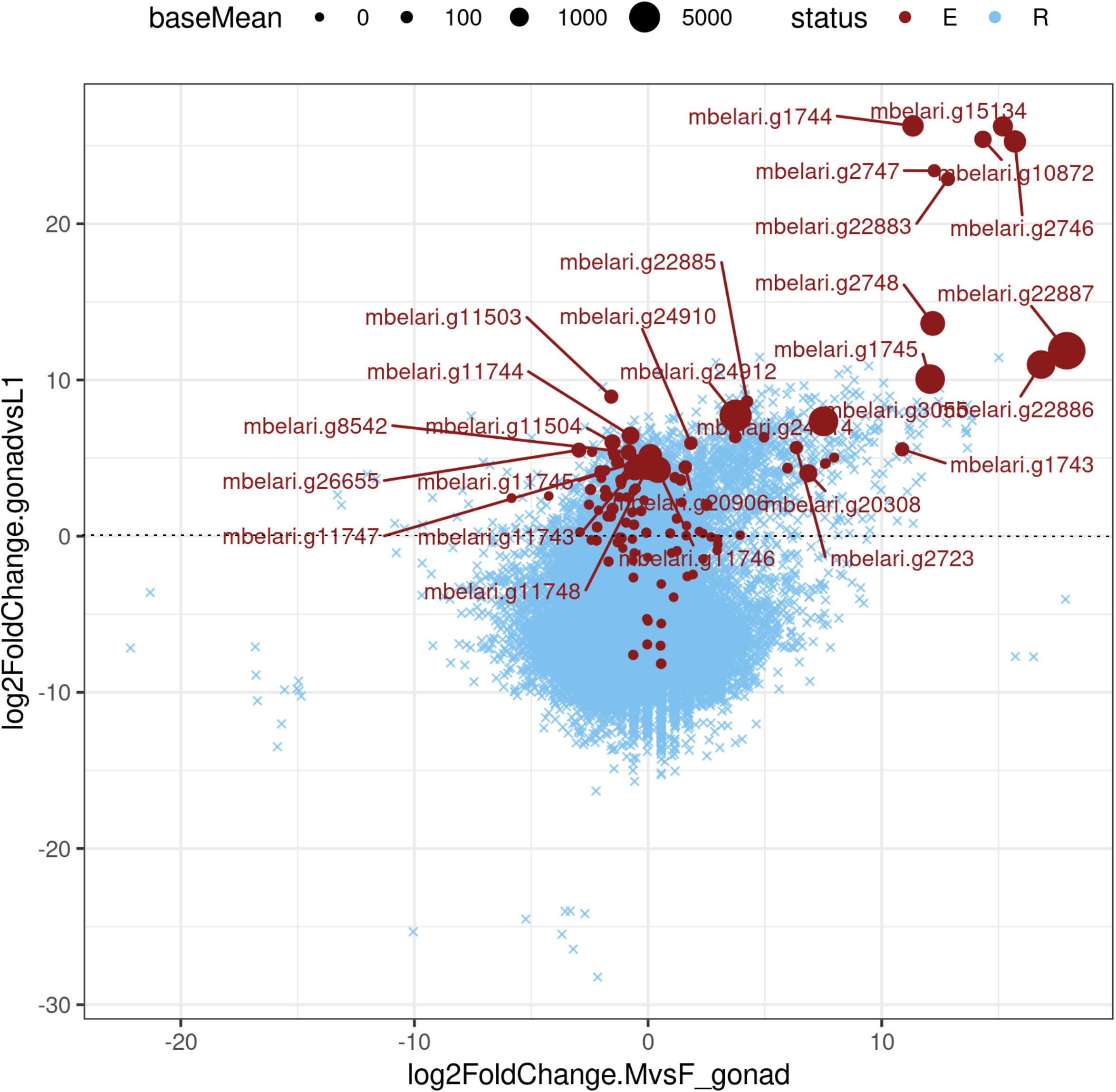
Analysis of the eliminated unique sequences in *M. belari*. Analysis of the RNAseq read counts between L1s and male and female gonads of *M. belari* (blue cross). The log2 fold change between read counts in gonads (both male and female) versus L1s is shown relative to the log2 fold change between male versus female gonads. Genes shown below the horizontal dotted black line are those specifically expressed in L1s, whereas those on the upper part are germline-specific expressed genes. Similarly, genes found on the left of the panel are female specific whereas those on the right are male specific. The eliminated genes whose base count is higher than 100 are shown on the graph with red dots. The biggest dots correspond to highly expressed genes. As expected, the genes eliminated in the soma are expressed only in the germline.

## Discussion

By serendipity, we discovered DNA fragmentation in fixed specimens of *M. belari* embryos. We found that these fragments appeared in the 5 presomatic blastomeres as they entered mitosis, and were absent in the P4 germline precursor cell, as previously described in *Ascaris* embryos. Our cytological observations combined with genome analysis revealed that similar to *Ascaris*, DNA elimination also leads to the removal of fragments of DNA. Similarly, we did not find evidence of chromosome fusion or recombination after elimination. Rather, the number of chromosomes increased after elimination suggesting that the retained fragments heal, most likely with the addition of new telomeres. Finally, elimination occurs independently in the 5 presomatic blastomeres. For each cell, elimination happens during a single mitosis, and not over several cell divisions.

In contrast to *Ascaris* however, we found that chromosomes are already fragmented in prometaphase of a mitosis of elimination. During metaphase, some fragments do not align on the metaphase plate, most likely because they have lost the kinetochore proteins. In *Ascaris*, metaphase spindles do not display any visible defects and fragmentation seems to occur during anaphase. This difference may simply reflect a difference in the progressive loss of kinetochore proteins between species. Alternatively, it is also possible that fragmentation occurs in interphase in *Mesorhabditis, for instance* because it depends on DNA replication, whereas another mechanism is responsible for fragmentation during anaphase in *Ascaris*.

As in other Metazoans undergoing PDE, we found that *Mesorhabditis* also eliminate satellite repeats, transposable elements and some protein-coding genes. Hence, in all species studied so far, PDE targets sequences that must be silenced in the soma (repeated elements and germline-specific genes). Therefore, PDE could have emerged as an efficient and irreversible mechanism to ensure chromatin silencing in the soma, by physically removing the unwanted sequences early in development rather than maintaining a costly silencing machinery throughout development. However, in Ascaris, in lampreys, and now in *Mesorhabditis*, although some germline specific genes are eliminated, they correspond to only a small fraction of all genes. Similarly, not all TE or satellite repeats are targeted by PDE, indicating that PDE does not fully substitute for the regular silencing machinery. Other evolutionary scenarios are possible (Drotos et al., 2022). The primary function of PDE may have been the silencing of some repeated elements and has been co-opted to eliminate more of those repeats or even genes that have a deleterious function in the soma. However, the elimination of germline genes may have evolved as a neutral trait (not having a selective advantage). Indeed, if PDE randomly targets protein coding genes, it will have no consequences on fitness only if those genes have no role in the soma. Hence, PDE may target germline specific genes only because those elimination are cryptic for selection. Our discovery that *Mesorhabditis* eliminate very few protein coding genes, which are in addition poorly conserved, is consistent with this latter scenario. Nevertheless, more work will be required to explore the exact function of the soma eliminated genes in *Mesorhabditis*, for instance the conserved sperm gene *spe-49*, and the consequences of their misexpression in the soma.

In a scenario where PDE is co-opted to eliminate genes in addition to repeated sequences, we expect that species in which PDE is very ancient eliminate more genes than species in which PDE has emerged recently. *Mesorhabditis* may belong to this category of species, for which PDE is a recent innovation. If true, *Mesorhabditis* species will constitute an interesting study system in which the mechanisms of PDE should be simpler to study, in contrast to the situation found in Ciliates where PDE, since its emergence, has become an extremely elaborate and complex system, with extensive diversification between species (Maurer-Alcalá and Nowacki, 2019; Rzeszutek et al., 2020).

150 years after the seminal discovery of PDE in *Ascaris* by Th. Boveri (Boveri, 1887), the mechanisms of PDE in animals are still mysterious. Our finding that a free-living nematode close to *C. elegans* undergoes PDE will open the way to functional approaches, and hopefully will contribute to answering some of the fascinating questions in the field. Besides having a small genome, these nematodes share the same experimental advantages as *C. elegans* (can be easily cultivated in the lab, can be frozen and synchronized), including the possibility to perform gene inactivation by CRISPR/Cas9 or RNAi (Blanc & al. in prep).

Finally, our fortuitous discovery of PDE in *Mesorhabditis* raises the exciting possibility that more systematic cytological and/or genomic exploration of Metazoans may uncover a larger number of species undergoing PDE than anticipated.

## Materials and methods

### Nematode maintenance

*Mesorhabditis* species are maintained at 20°C on NGM plates seeded with *E. coli* OP50, following *C. elegans* protocols (Grosmaire et al., 2019). For this study, we analyzed *M. belari* JU2817, *M. simplex* (JU2864), *M. spiculigera* AF72 (Launay et al., 2020), as well as *M. irregularis* (SB321) (a kind gift from David Fitch).

### *Estimation of germline/soma ratio in* M. belari

We fixed and stained L1 larvae and adults with Hoechst in order to obtain an estimation of the number of cells. Nuclei of the germline are easily recognizable, based on their round shape and organization (by analogy with *C. elegans*). Visual inspection and manual counting of the nuclei revealed that L1s have approximately 500 somatic nuclei and only one germ cell (as described in (Félix and Sternberg, 1996)). The gonad of adult females contained very few cells compared to *C. elegans*, with a maximum of 100 nuclei (also because they have a single gonadal arm, (Félix and Sternberg, 1996)), whereas males contained approximately 300 germline nuclei. It was difficult to estimate the number of somatic nuclei in adults and we assumed they had the same number as those found in *C. elegans* and other nematodes, i.e. ∼900 to 1000 somatic cells.

### DNA and RNA sample preparation for sequencing

We obtained a pure sample of L1 larvae, following protocols used for *C. elegans*. For DNA extraction, a pellet of larvae was snap frozen and 3 cycles of freezing and thawing were performed before adding 600 ul of Qiagen Cell lysis Solution and 6 ul of proteinase K (17mg/ml). After an overnight incubation at 65°C, we added 10 ul of RNAse A (10 mg/ul) for 1h at 37°C. Next, 200 ul of Qiagen Protein Precipitation Solution was added and samples were centrifuged for 10 min at 13000 rpm. From the supernatant, we performed DNA precipitation with isopropanol and the DNA pellet was next washed two times with EtOH 70%.

For RNA preparation, we used the SmartSeq2 protocol (Picelli et al., 2014). We manually selected single L1 larvae or dissected single gonads from male and female adults. We sequenced 5 samples per condition.

### Sequencing, data cleaning and trimming

DNA sequencing of adult males or females had previously been obtained (Grosmaire et al., 2019). Sequencing of DNA from L1s or cDNA from L1 and gonads was performed similarly on an Illumina HiSeq 4000 Sequencer with fragment length of 500bp and paired end reads of 100 bp.

We used fastp (v0.23.2, (Chen et al., 2018)) to clean data for bad quality reads (quality > 30). We also discarded reads coming from *E*.*coli* OP50 contamination using minimap2 (Li, 2018).

### K-mer analyses

We obtained similar results using different values of k. For simplicity, only the analysis performed with k=56 is presented in this study. K-mer were independently counted for each library using jellyfish (v2.3.0, (Marçais and Kingsford, 2011)). As the number of unique k-mer is huge, we created a subset by randomly drawing k-mer from a small genomic sequence (representing 1/500 of the genome) and kept the highest represented kmer for each library using *ad-hoc* scripts. K-mer showing interesting biased ratios were inspected and manually assembled to find repeated sequences. To normalize k-mer counts between DNA-seq libraries, k-mer raw counts were divided by the total number of counts per library and multiplied by the expected length of the genome (150Mb).

### Genome coverage

After cleaning, reads were mapped on the genome using minimap2 (Li, 2018). Then, alignment files (BAM) were converted into coverage for each 100 bp window of the genome using bedtools (Quinlan and Hall, 2010) and bedGraphToBigWig (Kent et al., 2010).

### Homology detection

We used Orthofinder ((Emms and Kelly, 2019), option: -S blast -M msa -I 0.8) to predict homology relationship between transcriptomes of *C. elegans, P. pacificus,O. tipulae*, and *M. belari* because few genes (< 30 %) have an available homology relationship predicted on https://parasite.wormbase.org/.

### Antibody and DNA stainings

Embryos were fixed with methanol and the freeze-cracking method, as described in (Launay et al., 2020). Embryos were stained with Hoechst (0.5 ug/ml). The following antibodies were used: mouse anti-tubulin (1/200, Sigma DM1A), secondary donkey anti-mouse Alexa 488 (1/1000, Jackson Immunoresearch, ref 115-545-166), secondary donkey anti-rabbit Cy3 (1/1000, Jackson Immunoresearch, ref 711-165-152). We also generated an antibody against *M. belari bub-1* (gene mbelari.g12204) (1/10000 Covalab). Two peptides C-ALNASKEKPEEQLD-coNH2 and C-SPIVEDQDHENSTNG-coNH2 were injected in rabbits and the purified serum obtained at day 74 was used for staining.

### DNA FISH

The following fluorescent oligo probes were ordered at Sigma and were used at 400 nM concentration (except the telomere probe which was used at 500 nM):

Telomeres: 5’ [TAM]TTAGGCTTAGGCTTAGGCTTAGGC 3’

*M. belari* Sat11: 5’ [6FAM]GGAACCGTATTGGAACCGTATTGGAACCGTATT 3’ or 5’ [TAM]GGAACCGTATTGGAACCGTATTGGAACCGTATT 3’

*M. belari* Sat65 5’[TAM]AGCTCAAATAGACTGAATAGCCTATTCTGAGCTGAATAGGCAATCTACACTTAATACCGTAATCC 3’

*M. spiculigera* Sat23 5’ [TAM]AGGTGCAGGAGCCGAAACTATTT 3’

Embryos and gonads were freeze-cracked and processed as described in (Grosmaire et al., 2019). Images were taken on an LSM-800 confocal.

### Soma-specific PCR

In order to validate the elimination of some sequences in the soma, we performed PCR on dissected nematode heads, which contain only somatic tissues (Figure S4). Worms were cut with a razor blade in a watch glass containing M9 and the head was pipetted and placed in 10 ul of Worm Lysis Buffer. The rest of the body, containing the germline, was also used as a control. Samples were incubated for 1h at 65°C followed by 10 min at 95°C and 2 ul of the mix was then used to perform a PCR with specific primers. We also systematically performed a PCR with Large Subunit rRNA (LSU) primers to confirm the presence of DNA in the tube LSU_foward 5’: acaagtaccgtgagggaaagttg, LSU_reverse 5’ :TCGGAAGGAACCAGCTACTA

## Supporting information

List of eliminated protein-coding genes

## Acknowledgments

We thank the French Institute of Bioinformatics—IFB CNRS UMS 3601—(funded as part of Investissement d’avenir program managed by Agence Nationale pour la Recherche, contract ANR-11-INBS-0013) for providing life science data and tools, storage and computing resources on the IFB national service infrastructure in bioinformatics. We acknowledge the contribution of the imaging platform PLATIM from SFR Biosciences Lyon, as well as the sequencing platform Genomeast_ Strasbourg. We thank Clara Bourgeais, Ingrid Dourlens, Remy Mimbre and Caroline Blanc for technical help and Michael Ailion for critical reading of the manuscript.

## Supplementary Material

**Table S1: list of eliminated protein-coding genes** and RNAseq read counts in the entire population, in L1s, in male gonads and female gonads.

**Figure S1:**
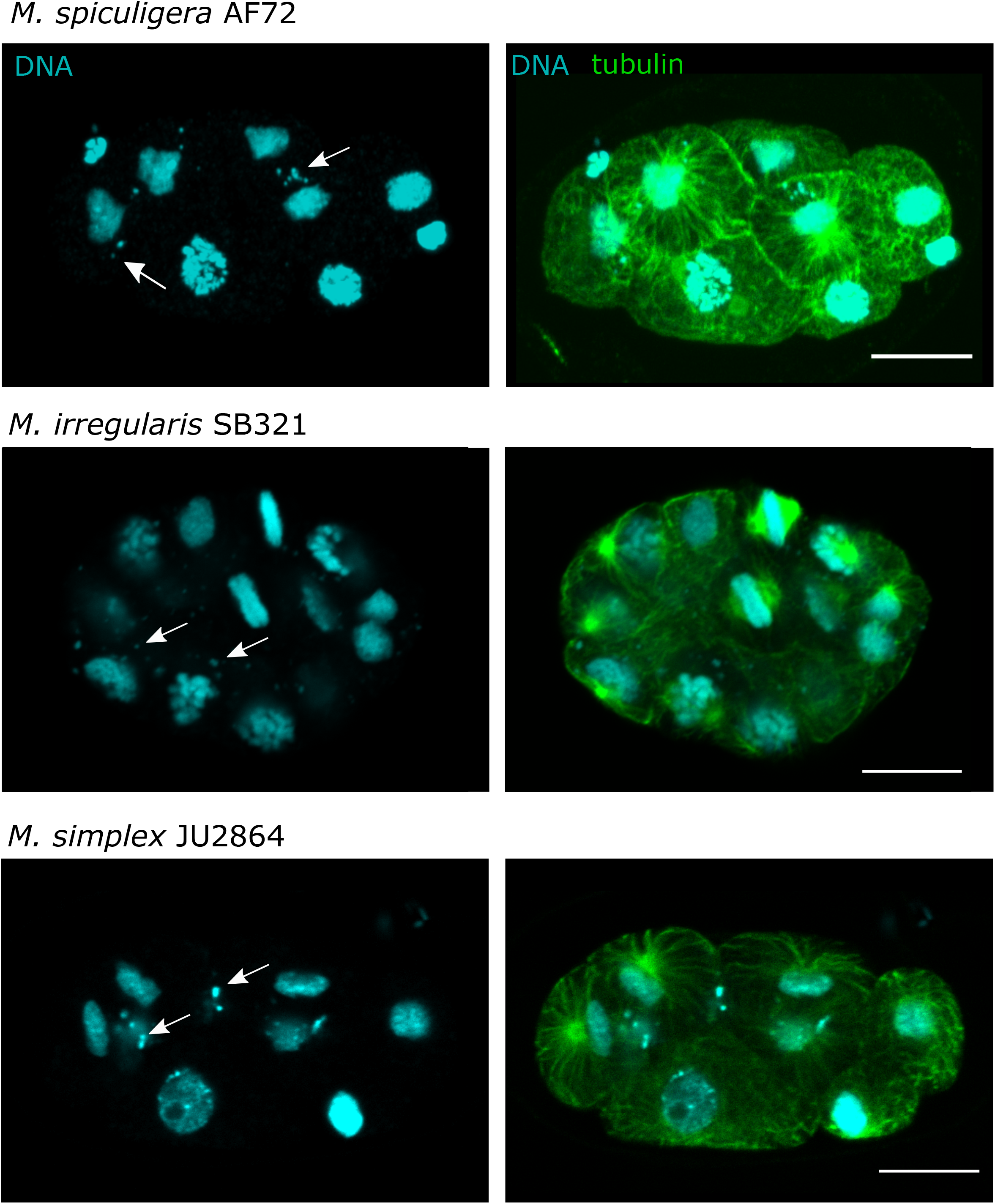
Cytological evidence of DNA elimination in other *Mesorhabditis* species. One representative embryo per species is shown for the sexual species *M. spiculigera* and *M. irregularis* and the asexual species *M. simplex*. Tubulin is in green, DNA in blue. Arrows show the unretained DNA fragments. Scale bar is 10 um.

**Figure S2:**
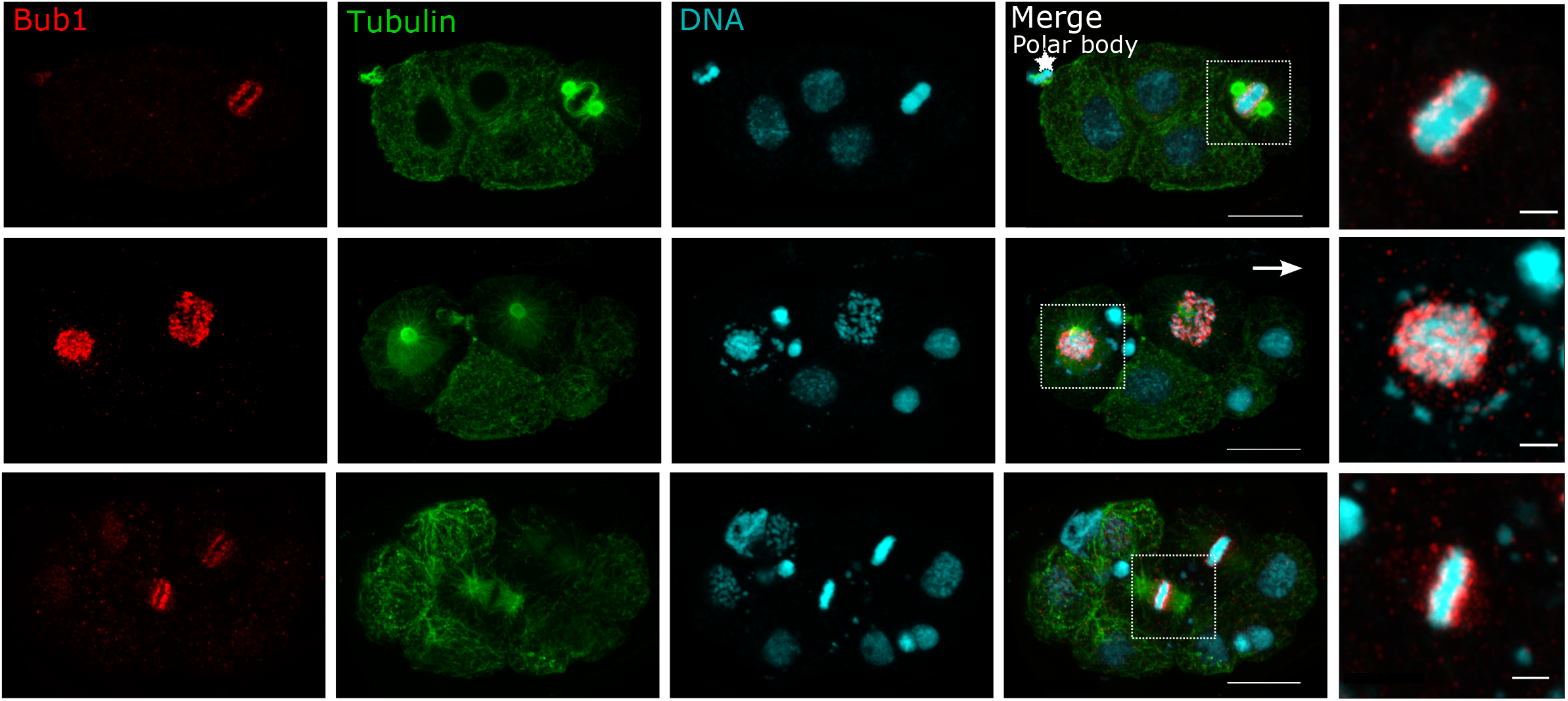
BUB-1 localisation in *M. belari* embryos. Immunostaining of embryos with an anti-BUB-1 antibody, which labels kinetochores. *M. belari* are holocentric, hence BUB-1 is uniformly distributed along chromosomes during mitosis, as shown in the top embryo before a division of elimination. The middle and bottom embryos are shown during a division of elimination and insets are shown on the right. BUB-1 is found only on the retained fragments (metaphase plate) and not on the halo of unretained fragments. Tubulin is in green, BUB-1 is in red, and DNA is in blue. Scale bar is 10 and 2 um for the embryos or insets, respectively.

**Figure S3:**
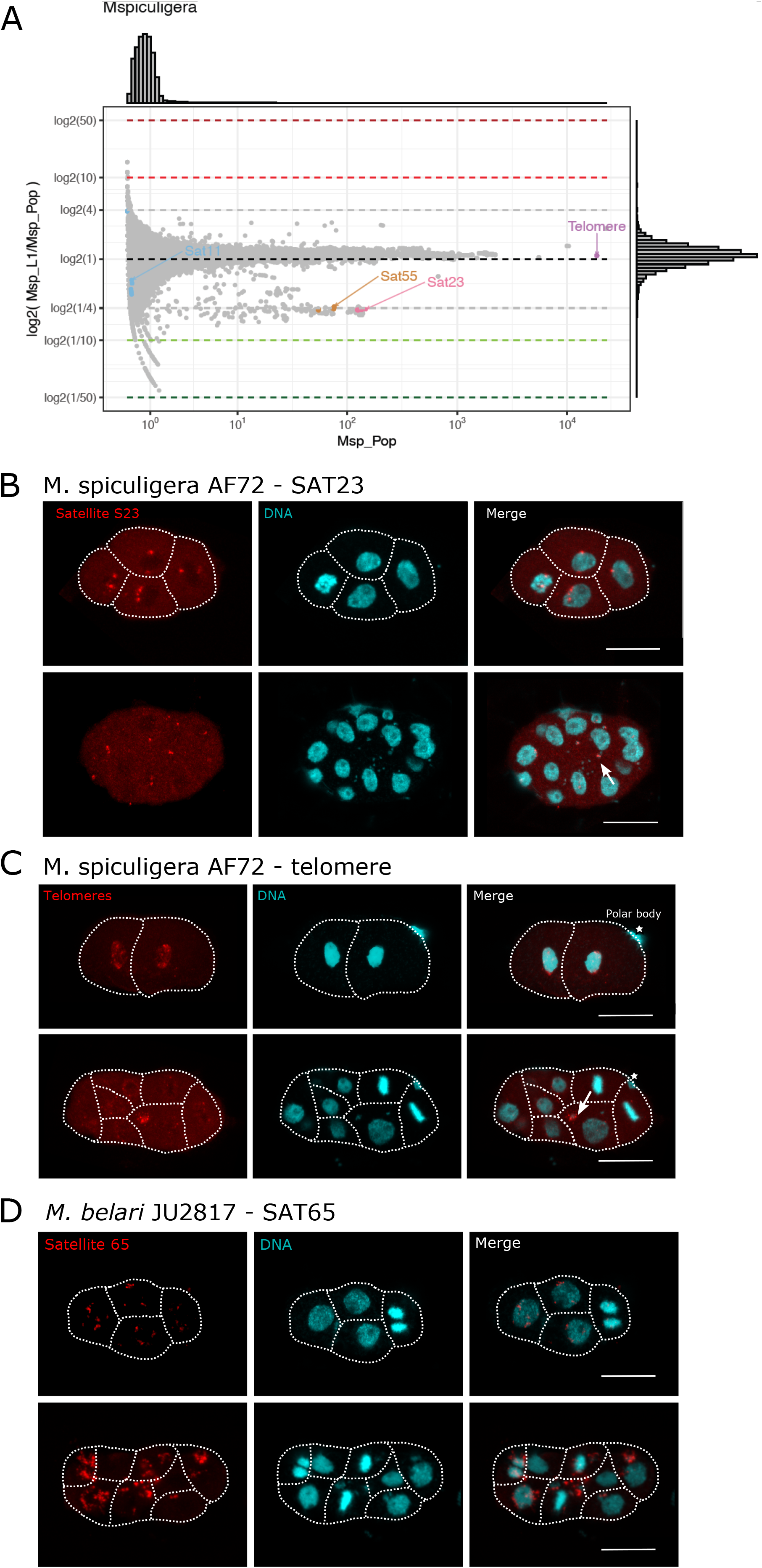
Identification of the eliminated satellite-tandem repeats in *M. belari* and *M. spiculigera*. (A). k-mer analysis in *M. spiculigera* as shown in Figure 3 for *M. belari*. The ratio of counts between L1s and males is shown in log2 relative to the counts found in males. One grey dot represents one 56-mer. The majority of k-mers are unique and equally represented in both samples (log2(1)). K-mers containing the telomeric repeat are overrepresented in males and are overall extremely abundant (5.10e4). Some k-mers, highlighted in colors, are underrepresented in L1s. K-mers shown in orange and pink are different versions of the same 55nt and 23nt sequences, respectively. (B). DNA FISH validation of *M. spiculigera*_Sat23 elimination (in red) before and during elimination. (C). DNA FISH validation of *M. spiculigera*_telomere elimination (in red) before and during elimination. Note that the P4 cell is not shown in this focal plane. (D). DNA FISH validation of *M. belari*_Sat65 elimination (in red) before (upper embryo) and during elimination. The white stars show the polar body. DNA is in blue. Scale bar is 10 uM.

**Figure S4:**
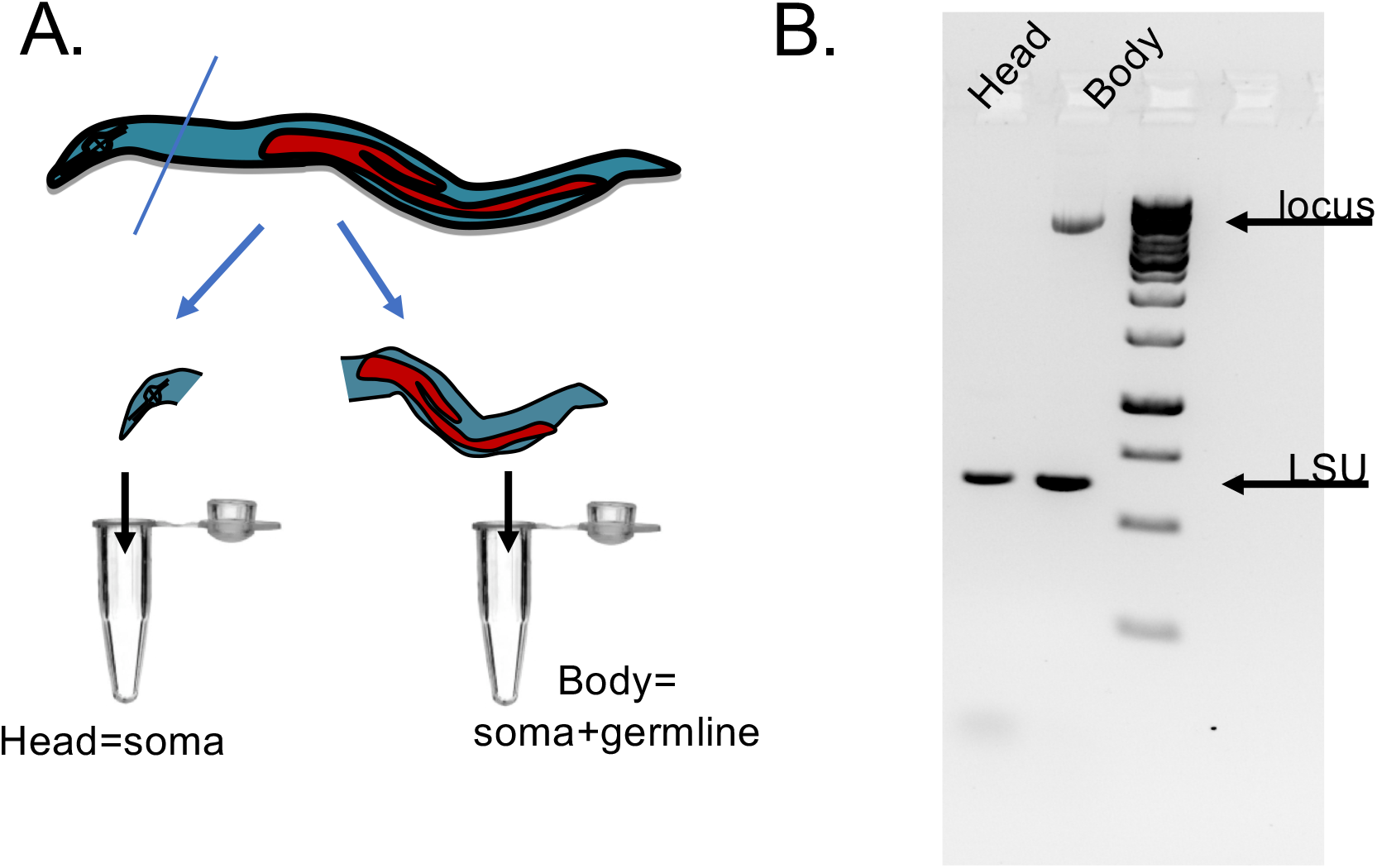
Soma-specific PCR. (A).Schematic representation of soma-specific PCR. Adults are cut and the head, which contains only somatic tissues, is isolated from the rest of the body, which contains soma and germline tissues. (B). PCRs are performed on the head or body part of a single worm. Example of a locus which is eliminated in the soma. Primers 5’ mbelari_1060_Fw :5’-ACCTGTACACCATTGAGACTGA and primer mbelari_1060_Rv :5’-ACTGCGTTCGCCGTACAA were used as well as primers to amplify the LSU locus which is not eliminated.

## References

Boveri, T. (1887). Ueber Differenzierung der Zellkerne wahrend der Furchung des Eies von Ascaris megalocephala. Anat Anz 2, 688–693.

Bracht, J.R., Fang, W., Goldman, A.D., Dolzhenko, E., Stein, E.M., and Landweber, L.F. (2013). Genomes on the edge: programmed genome instability in ciliates. Cell 152, 406–416.

Chen, S., Zhou, Y., Chen, Y., and Gu, J. (2018). Fastp: An Ultra-Fast All-in-One FASTQ Preprocessor. Bioinformatics 34, i884–i890.

Dedukh, D., and Krasikova, A. (2022). Delete and survive: strategies of programmed genetic material elimination in eukaryotes. Biol. Rev. Camb. Philos. Soc. 97, 195–216.

Drotos, K.H.I., Zagoskin, M.V., Kess, T., Gregory, T.R., and Wyngaard, G.A. (2022). Throwing away DNA: programmed downsizing in somatic nuclei. Trends Genet. 0.

Emms, D.M., and Kelly, S. (2019). OrthoFinder: phylogenetic orthology inference for comparative genomics. Genome Biol. 20, 238.

Félix, M.-A., and Sternberg, P.W. (1996). Symmetry breakage in the development of one-armed gonads in nematodes. Development 122, 2129–2142.

Gonzalez de la Rosa, P.M., Thomson, M., Trivedi, U., Tracey, A., Tandonnet, S., and Blaxter, M. (2021). A telomere-to-telomere assembly of Oscheius tipulae and the evolution of rhabditid nematode chromosomes. G3 Bethesda Md 11, jkaa020.

Grishanin, A. (2014). Chromatin diminution in Copepoda (Crustacea): pattern, biological role and evolutionary aspects. Comp. Cytogenet. 8, 1–10.

Grosmaire, M., Launay, C., Siegwald, M., Brugière, T., Estrada-Virrueta, L., Berger, D., Burny, C., Modolo, L., Blaxter, M., Meister, P., et al. (2019). Males as somatic investment in a parthenogenetic nematode. Science 363, 1210–1213.

Haag, E.S., Fitch, D.H.A., and Delattre, M. (2018). From “the Worm” to “the Worms” and Back Again: The Evolutionary Developmental Biology of Nematodes. Genetics 210, 397–433.

Kang, Y., Wang, J., Neff, A., Kratzer, S., Kimura, H., and Davis, R.E. (2016). Differential Chromosomal Localization of Centromeric Histone CENP-A Contributes to Nematode Programmed DNA Elimination. Cell Rep. 16, 2308–2316.

Kent, W.J., Zweig, A.S., Barber, G., Hinrichs, A.S., and Karolchik, D. (2010). BigWig and BigBed: enabling browsing of large distributed datasets. Bioinformatics 26, 2204– 2207.

Launay, C., Félix, M.-A., Dieng, J., and Delattre, M. (2020). Diversification and hybrid incompatibility in auto-pseudogamous species of Mesorhabditis nematodes. BMC Evol. Biol. 20, 105.

Li, H. (2018). Minimap2: Pairwise Alignment for Nucleotide Sequences. Bioinformatics 34, 3094–3100.

Marçais, G., and Kingsford, C. (2011). A fast, lock-free approach for efficient parallel counting of occurrences of k-mers. Bioinformatics 27, 764–770.

Maurer-Alcalá, X.X., and Nowacki, M. (2019). Evolutionary origins and impacts of genome architecture in ciliates. Ann. N. Y. Acad. Sci. 1447, 110–118.

Niedermaier, J., and Moritz, K.B. (2000). Organization and dynamics of satellite and telomere DNAs in Ascaris: implications for formation and programmed breakdown of compound chromosomes. Chromosoma 109, 439–452.

Picelli, S., Faridani, O.R., Björklund, Å.K., Winberg, G., Sagasser, S., and Sandberg, R. (2014). Full-length RNA-seq from single cells using Smart-seq2. Nat. Protoc. 9, 171–181.

Quinlan, A.R., and Hall, I.M. (2010). BEDTools: a flexible suite of utilities for comparing genomic features. Bioinformatics 26, 841–842.

Rzeszutek, I., Maurer-Alcalá, X.X., and Nowacki, M. (2020). Programmed genome rearrangements in ciliates. Cell. Mol. Life Sci. CMLS 77, 4615–4629.

Smith, J.J., Timoshevskaya, N., Ye, C., Holt, C., Keinath, M.C., Parker, H.J., Cook, M.E., Hess, J.E., Narum, S.R., Lamanna, F., et al. (2018). The sea lamprey germline genome provides insights into programmed genome rearrangement and vertebrate evolution. Nat. Genet. 50, 270–277.

Timoshevskiy, V.A., Herdy, J.R., Keinath, M.C., and Smith, J.J. (2016). Cellular and Molecular Features of Developmentally Programmed Genome Rearrangement in a Vertebrate (Sea Lamprey: Petromyzon marinus). PLoS Genet. 12, e1006103.

Torgasheva, A.A., Malinovskaya, L.P., Zadesenets, K.S., Karamysheva, T.V., Kizilova, E.A., Akberdina, E.A., Pristyazhnyuk, I.E., Shnaider, E.P., Volodkina, V.A., Saifitdinova, A.F., et al. (2019). Germline-restricted chromosome (GRC) is widespread among songbirds. Proc. Natl. Acad. Sci. U. S. A. 116, 11845–11850.

Wang, J., and Davis, R.E. (2014). Programmed DNA elimination in multicellular organisms. Curr. Opin. Genet. Dev. 27, 26–34.

Wang, J., Mitreva, M., Berriman, M., Thorne, A., Magrini, V., Koutsovoulos, G., Kumar, S., Blaxter, M.L., and Davis, R.E. (2012). Silencing of germline-expressed genes by DNA elimination in somatic cells. Dev. Cell 23, 1072–1080.

Wang, J., Gao, S., Mostovoy, Y., Kang, Y., Zagoskin, M., Sun, Y., Zhang, B., White, L.K., Easton, A., Nutman, T.B., et al. (2017). Comparative genome analysis of programmed DNA elimination in nematodes. Genome Res. 27, 2001–2014.

Wang, J., Veronezi, G.M.B., Kang, Y., Zagoskin, M., O’Toole, E.T., and Davis, R.E. (2020). Comprehensive Chromosome End Remodeling during Programmed DNA Elimination. Curr. Biol. CB 30, 3397-3413.e4.

